# Biomarker signatures of quality for nasal chondrocyte-derived engineered cartilage

**DOI:** 10.1101/2020.02.12.945527

**Authors:** M. Adelaide Asnaghi, Laura Power, Andrea Barbero, Martin Haug, Ruth Köppl, David Wendt, Ivan Martin

**Author notes:** Both authors contributed equally (M. Adelaide Asnaghi, Laura Power). **Correspondence:** Prof. Ivan Martin.

## Abstract

The definition of quality controls for cell therapy and engineered product manufacturing processes is critical for safe, effective, and standardized clinical implementation. Using the example context of cartilage grafts engineered from autologous nasal chondrocytes, currently used for articular cartilage repair in a phase II clinical trial, we outlined how gene expression patterns and generalized linear models can be introduced to define molecular signatures of identity, purity, and potency. We first verified that cells from the biopsied nasal cartilage can be contaminated by cells from a neighboring tissue, namely perichondrial cells, and discovered that they cannot deposit cartilaginous matrix. Differential analysis of gene expression enabled the definition of identity markers for the two cell populations, which were predictive of purity in mixed cultures. Specific patterns of expression of the same genes were significantly correlated with cell potency, defined as the capacity to generate tissues with histological and biochemical features of hyaline cartilage. The outlined approach can now be considered for implementation in a Good Manufacturing Practice setting, and offers a paradigm for other regenerative cellular therapies.

## 1 Introduction

Large cartilage defects in adults have limited capacity to regenerate, and state-of-the art regenerative medicine therapies do not induce reproducible or stable results (Teo et al., 2019; Kon et al., 2009). We previously demonstrated the safety and feasibility of autologous nasal chondrocyte-derived engineered cartilage for the treatment of focal traumatic lesions in the knee in a phase I clinical trial (Mumme et al., 2016), and a phase II clinical trial is ongoing to investigate efficacy. Briefly, autologous nasal chondrocytes are expanded in vitro before seeding onto a collagen I/III scaffold and cultured in chondrogenic conditions to produce a mature, hyaline-like cartilage graft that is then implanted into the knee cartilage defect of the same patient.

The starting material for this approach is a biopsy from the native nasal septum cartilage, which, like articular cartilage, is a hyaline cartilage (Al Dayeh et al., 2013) composed predominantly of water, type II collagen, glycosaminoglycan (GAG) containing proteoglycans, and the one cell type, chondrocytes (Buckwalter et al., 1998). Mucoperichondrium is the tissue that overlays nasal cartilage; it consists of several layers, including mucosa, lamina propria, and perichondrium, the tissue directly adjacent to the cartilage that is tightly attached and cannot be easily distinguished (Aksoy et al., 2012). Currently, nasal septal cartilage and mucoperichondrium are separated by pulling them apart with forceps and are identified based on their physical characteristics. Due to donor-related and operator-related variability, the resulting biopsy may not be completely pure after cleaning, with some overlaying tissue remaining attached to the cartilage.

When working with intrinsically variable donor-derived human materials, such as tissues and cells, establishing the quality and consistency of the starting material is key to ensuring reproducibly high quality engineered products, not least to avoid the consequences of cell misidentification (Hyun-woo, 2019; Editorial, 2009). This prompts for the development of identity and purity assays (Carmen et al., 2012), which can be based on various characteristics of the cells, such as gene or protein expression. Gene expression markers have been proposed for articular cartilage cell identity and purity assays (CHMP 2017; Diaz-Romero et al., 2017; Bravery et al., 2013; Mollenhauer et al 2010). However, until now, there are no known biomarkers that can distinguish the cell types found in nasal cartilage biopsies. Moreover, the impact of possibly contaminating cells on nasal chondrocyte-based engineered cartilage has not been investigated.

Engineered products treat diseases or damage through repairing, replacing, or regenerating tissues or organs (Detela et al., 2019). A potency assay must be developed based on the mode of action of the tissue engineered product (CHMP 2008)–in our case, the filling of cartilage defects with healthy, hyaline-like tissue–which is ideally correlated to the efficacy, leading to consistent quality of the tissue engineered product and good clinical outcome (Bravery et al., 2013).

In this study, we first investigate which cell types are potentially contaminating in human nasal septum cartilage biopsies and their impact on the quality of engineered cartilage. We then investigated whether gene expression analysis could discriminate the contaminant cells found in nasal septal biopsies for the development of a characterization panel for identity and purity quality controls. To assess potency, we compared the gene expression of nasal septum biopsy-derived cells to their ability to produce cartilaginous tissue. Finally, we propose how these purity and potency assays could be implemented in a Good Manufacturing Practice (GMP) compliant process for the translation of our regenerative therapy product.

## 2 Materials and methods

### 2.1 Cell isolation and expansion

Human nasal septum biopsies were collected from 17 donors (9 female, 8 male, mean age 46 years, range 16–84 years) after informed consent and in accordance with the local ethical commission (EKBB; Ref.# 78/07). Two samples derived from patients enrolled in the Nose2Knee clinical trials (ClinicalTrials.gov, number NCT01605201 and number NCT02673905).

For 4 donors, the biopsy was carefully dissected to give a pure cartilage sample (NC) and a pure perichondrium sample (PC). Nasal chondrocytes were isolated from NC by enzymatic digestion as previously described (Jakob et al., 2003) with 0.15% collagenase II (Worthington) for 22 hours at 37°C. After digestion, cells were plated in tissue culture flasks at a density of 1 × 10^4^ cells/cm^2^ and cultured in medium consisting of *complete medium* (Dulbecco’s Modified Eagle’s Medium (DMEM) containing 4.5 mg/mL D-glucose and 0.1 mM nonessential amino acids, 10% fetal bovine serum (FBS), 1 mM sodium pyruvate, 100 mM HEPES buffer, 100 U/mL penicillin, 100 μg/mL streptomycin, and 0.29 mg/mL L-glutamine (all from Invitrogen), supplemented with 1 ng/mL transforming growth factor beta-1 (TGF-β1) and 5 ng/mL fibroblast growth factor-2 (FGF-2) (both from R&D Systems) at 37°C and 5% CO2 in a humidified incubator (Thermo Scientific Heraeus) as previously described (Jakob et al., 2003). When approaching 80% confluence, cells were detached using 0.05% trypsin-EDTA (Invitrogen) and re-plated.

PC tissue samples were cut in small pieces and put on the bottom of plastic culture dishes to isolate adherent cells that migrated out of the tissue for one week in complete medium. Cells were then detached using 0.05% trypsin-EDTA and further cultured until confluence in the same conditions as nasal chondrocytes.

Specific ratios of NC to PC cells were combined at passage two to generate mixed populations of known purity of 100%, 90%, 80%, 70%, 60%, and 0% NC.

For all other biopsies, in case perichondrium was present, half of the sample was dissected removing all perichondrium to obtain a pure nasal cartilage sample (NC), while the overlaying perichondrium remained intact on the other half (NC+PC samples). Cells were isolated from each tissue sample by enzymatic digestion and expanded in complete medium supplemented with TGF-β1 and FGF-2 up to 2 passages as described above for nasal chondrocytes.

### 2.2 Chondrogenic redifferentiation

#### 2.2.1 Micromass pellets

Cells expanded until passage two were redifferentiated by culturing as 3D micromass pellets, as previously described (Asnaghi et al., 2018). 3D micromass pellets were formed by centrifuging 5×10^5^ cells at 300 x *g* in 1.5 mL conical tubes (Sarstedt) and cultured for two weeks in chondrogenic serum-free medium consisting of DMEM containing 1 mM sodium pyruvate, 100 mM HEPES buffer, 100 U/mL penicillin, 100 μg/mL streptomycin, 0.29 g/mL L-glutamine, 1.25 mg/mL human serum albumin (CSL Behring), and 100nM dexamethasone (Sigma, Switzerland), supplemented with 10 ng/mL TGF-β1 (R&D), ITS+1 (10 µg/mL insulin, 5.5 µg/mL transferrin, 5 ng/mL selenium; Gibco), 100 µM ascorbic acid 2-phosphate (Sigma), and 4.7 µg/mL linoleic acid (Sigma). Culture medium was changed twice weekly.

#### 2.2.2 Engineered cartilage on Chondro-Gide

Passage two cells were seeded on collagen type I/III membranes (Chondro-Gide; Geistlich Pharma AG) at a density of 4.17 million cells per cm^2^. The resulting constructs were cultured for two weeks in chondrogenic medium consisting of complete medium supplemented with 10 μg/mL insulin (Novo Nordisk), and 0.1 mM ascorbic acid 2-phosphate (Sigma) at 37°C and 5% CO2 with media changes twice/week.

The described protocols match the ones used in the context of the clinical trial, where GMP-grade reagents and autologous serum instead of FBS are used. Grafts for clinical use are produced at the GMP facility at the University Hospital Basel according to standard operating procedures under a quality management system, as described in Mumme *et al* (Mumme et al., 2016).

### 2.3 Histology and immunohistochemistry

Samples were fixed overnight in 4% formalin and embedded in paraffin. Sections 5 µm in thickness were stained with safranin-O for glycosaminoglycans (GAG) and hematoxylin as a nuclear counterstaining as described elsewhere (Grogan et al., 2016). Immunohistochemistry against collagen type I (No. 0863170, MPBiomedicals, 1:5000) and collagen type II (No. 0863171, MPBiomedicals, 1:1000) was performed using the Vectastain ABC kit (Vector Labs) with hematoxylin counterstaining as in standard protocols (Scotti et al., 2010). Immunofluorescence staining against HAPLN1 was performed using the HAPLN1 primary antibody (ABIN653748, Abgent) after epitope heat retrieval at 95°C for 60 min and permeabillization for 5 min with 0.5% Triton-X, and a goat anti-rabbit Alexa Fluor 488 secondary antibody (Invitrogen, 1:200) with DAPI (Thermo Fisher Scientific D1306, 300mM) as a nuclear counterstain. Fluorescence images were acquired with a confocal microscope (Zeiss LSM 710).

Histological scoring via the Modified Bern Score (MBS) was performed on safranin O-stained histological images as previously described (Lehoczky et al., 2019; Grogan et al., 2006). Briefly, the MBS has two rating parameters that each receive a score between 0 and 3. First, the intensity of safranin-O staining (0 = no stain; 1 = weak staining; 2 = moderately even staining; 3 = even dark stain), and second, the morphology of the cells (0 = condensed/necrotic/pycnotic bodies; 1 = spindle/fibrous; 2 = mixed spindle/fibrous with rounded chondrogenic morphology; 3 = majority rounded/chondrogenic). The two values are summed together resulting in a maximum possible MBS of 6.

### 2.4 Quantitative qPCR

We chose the gene expression markers to investigate based on a literature search. The gene expression ratios of collagen II to I and aggrecan to versican are well-known chondrogenic markers (Martin et al., 2001). HAPLN1 has been found in most types of cartilage (Spicer et al., 2003), including in bovine nasal cartilage (Baker et al., 1978). Versican protein expression has been found in perichondrium from other cartilage tissue sources (Shibata et al., 2014) and nestin has been shown to be expressed in embryonic perichondrium (Ono et al., 2014). MFAP5 is found in elastic as well as non-elastic extracellular matrixes (Halper et al., 2014) and has been used as a negative marker for chondrogenic cells from articular cartilage (Rapko et al., 2010).

Total RNA was extracted from expanded cells at both P1 and P2, 3D micromass pellets, and engineered cartilage grafts with the Quick RNA Miniprep Plus kit (Zymo Research) and quantitative gene expression analysis was performed as previously described (Martin et al., 2001). Reverse transcription into cDNA was done from 3 μg of RNA by using 500 μg/mL random hexamers (Promega, Switzerland) and 0.5 μL of 200 UI/mL SuperScript III reverse transcriptase (Invitrogen). Assay on demand was used with TaqMan Gene Expression Master Mix to amplify type I collagen (Col I, Hs00164004), type II collagen (Col II, Hs00264051), aggrecan (Agg, Hs00153936_m1), Versican (Ver, Hs00171642_m1), link protein 1 (HAPLN1, Hs00157103_m1), MFAP5 (MFAP5, Hs00185803_m1), and nestin (Nes, Hs00707120_s1) (all from Applied Biosystems). The threshold cycle (CT) value of the reference gene, GAPDH (GAPDH, Hs00233992_m1), was subtracted from the CT value of the gene of interest to derive ΔC_T_ values. All displayed gene expression levels are, and statistical analyses were performed on, the ΔC_T_ values. GAPDH was found to be a stable reference gene for perichondrial cells.

### 2.5 Biochemical quantification of GAG and DNA

Samples of engineered cartilage and micromass pellets were digested with proteinase K (1 mg/mL proteinase K in 50 mM Tris with 1 mM EDTA, 1 mM iodoacetamide, and 10 mg/mL pepstatin A). The GAG content was determined by spectrophotometry using Dimethylmethylene Blue (Sigma-Aldrich 341088) as previously described (Barbosa et al., 2003). DNA content was measured using the CyQuant cell proliferation assay kit (Invitrogen).

### 2.6 Modeling

The generalized linear modeling (glm) function in R was used to build all the models. A logistic regression model was used to predict purity, where the response is a continuous probability between 0 (pure perichondrium) and 1 (pure cartilage) with samples from four donors and 48 independent experiments of known purities. For the logistic regression models, the McFadden pseudo R^2^ values were calculated with the pscl R package (Jackman et al., 2017) and the Hosmer-Lemeshow analysis was performed with the ResourceSelection R package (Lele et al., 2019). For the potency assay predicting GAG production, a gamma GLM with a log link was used to model quantified amounts of GAG (measured in µg). The MBS of chondrogenic pellets was modeled by first dividing the value by six, the maximum possible score, then training a multiple logistic regression model; the predicted responses were then multiplied by six. Samples from nine donors in 28 independent experiments were used to train the MBS potency assay and 25 independent experiments were used to train the GAG potency model and for gene selection. For all three assays, stepwise selection (Agostini et al., 2015) was performed in both directions; collagen II and I, aggrecan, versican, HAPLN1, and MFAP5 were tested and the model with the lowest Akaike information criterion (AIC) was chosen. Samples from five donors in 12 independent experiments were used to test the potency models. Residual plots were used to verify all the models. The correlation between the predicted and actual purity, GAG, and MBS values were calculated with the square of the Pearson correlation coefficient. The final equations of the potency models were rebuilt with both the training and test data together.

### 2.7 Statistical analysis

All calculations were performed using standard functions, unless otherwise stated, in R (R Core Team 2019). Statistical significance is defined as p < 0.05. Statistical significance for comparing two means was calculated using paired or unpaired t-tests and normality was checked with the Shapiro-Wilks test. To test multiple comparisons, a linear model was fitted, then the glht function of the multcomp R package (Hothorn et al., 2008) was used to test all the contrasts; p-values were corrected for multiple testing using the single-step Bonferroni method. Correlation plots using Spearman correlation coefficients (ρ) were created with the corrplot R package (Wei et al., 2017). Data are presented as mean and standard deviation of independent experiments with cells from at least 4 different donors. For each analysis at least 2 replicate micromass pellets were used per condition. Symbols used are: *** p<0.001, ** p<0.01, * p<0.05, and. p<0.1.

## 3 Results

### 3.1 Native nasal septum biopsy characterization

In the context of ongoing clinical trials, the nasal septum biopsy is harvested along the subperichondrial axis, so that most of the perichondrium remains in place in the patient’s nose, not only an efficient risk-control measure, but also important for the stability and healing of the donor site. More heterogeneous samples are obtained from plastic surgeries unrelated to clinical trials, which include mixed cartilage and perichondrium. Safranin O staining of nasal septum specimens indicated the presence of tissues with distinct characteristics, i.e., GAG-rich cartilage with round chondrocytes residing in lacunae and adjacent GAG-negative perichondrium containing cells with fibroblast-like morphology, comparable to previous findings (Bairati et al., 1996). Immunohistochemical analysis showed more collagen II in the cartilage and more collagen I in the perichondrium, confirming previously reported results (Popko et al., 2007). The border between the two tissues is not clearly defined in our samples, as in previous reports (Bleys et al., 2007; Bairati et al., 1996; Fig 1A).

**Figure 1.**
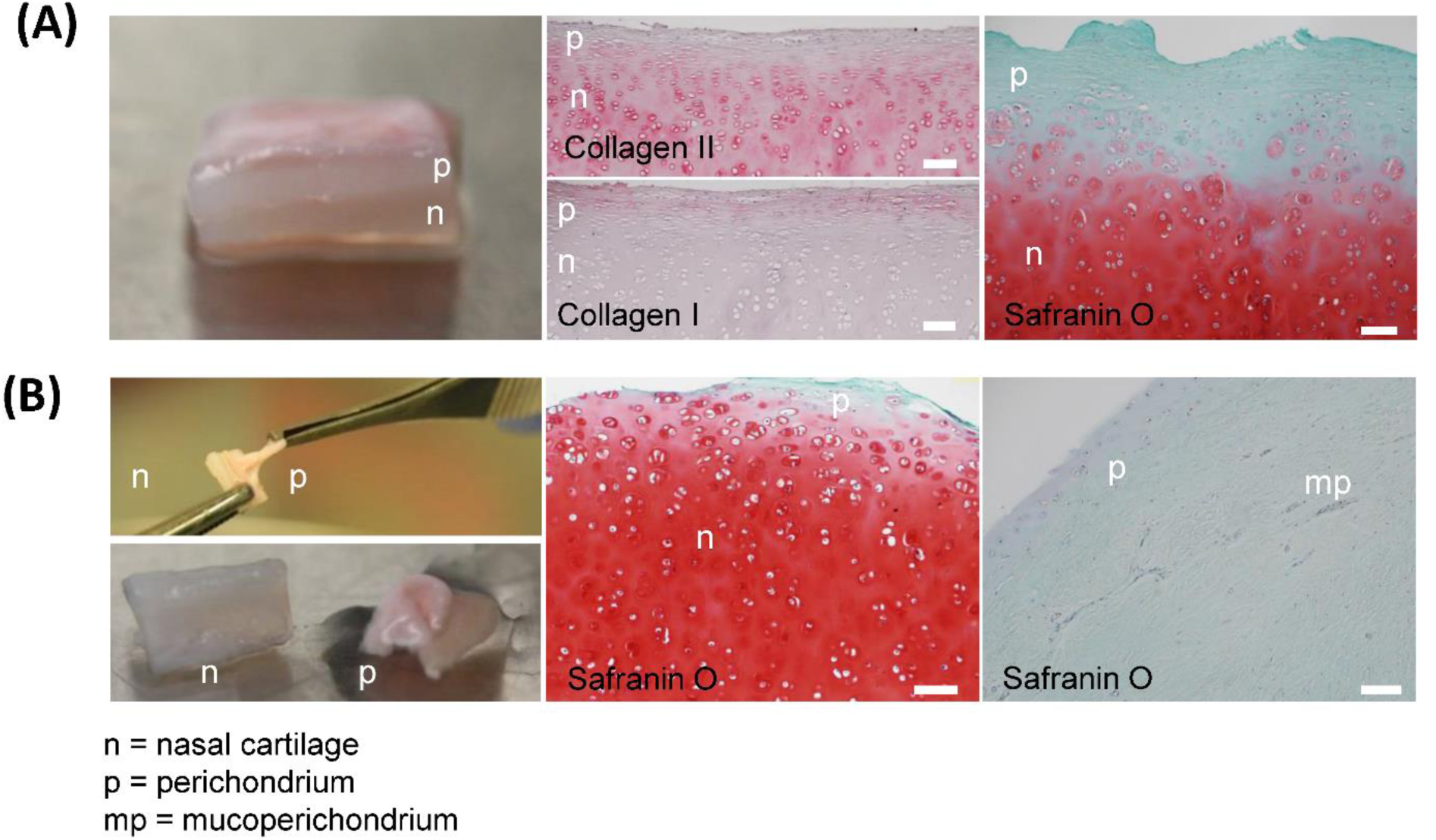
Native nasal septal cartilage and overlaying tissue. **(A)** Photograph of native nasal cartilage with overlaying tissues. Safranin O-stained histological image, HAPLN1 immunofluorescence, and collagen I, collagen II, and versican immunohistochemical images. **(B)** Photograph of the forceps separation technique and resulting separate nasal cartilage (n) and perichondrial tissue (p). Safranin O-stained histological images of separated nasal cartilage and perichondrial tissues. Scale bars are 100 µm

The separation of the cartilage and overlaying tissue is done by pulling them apart with forceps; however, the efficiency of this technique is unknown. Histological analysis after physical separation of cartilage and perichondrium revealed that the resulting biopsy may have small amounts of safranin O-negative tissue on the cartilage after separation (Fig 1B). This safranin O-negative region includes cambium, which is hypothesized to be the source of cells with tissue forming capacity (Van Osch et al., 2000; Upton et al., 1981), and sometimes perichondrium that is difficult to remove (Hellingman et al., 2011). Deeper cleaning of the starting biopsy (e.g., via scraping or cutting with a scalpel) is not a suitable option, since we observed reduced cell yield and slightly lower chondrogenic capacity in preliminary experiments, supporting the theory that this superficial region contains more potent cells.

### 3.2 Characterization of perichondrial cells

The samples we classify as pure nasal cartilage (NC) and perichondrium (PC) are the tissues after separation using the aforementioned technique. Visually, under macroscopic observation during expansion in cell culture dishes, NC and PC cells are not distinguishable, both having the same characteristic fibroblastic-like cell morphology. The proliferation rates of the two cell types were measured and found to be about equal (Fig 2A). To compare the chondrogenic capacity of NC and PC cells, we engineered pellets and found that NCs could reproducibly produce GAG and collagen II while PCs could not and predominantly produced type I collagen, as seen by histological analyses and biochemical quantification (Fig 2B,C).

**Figure 2.**
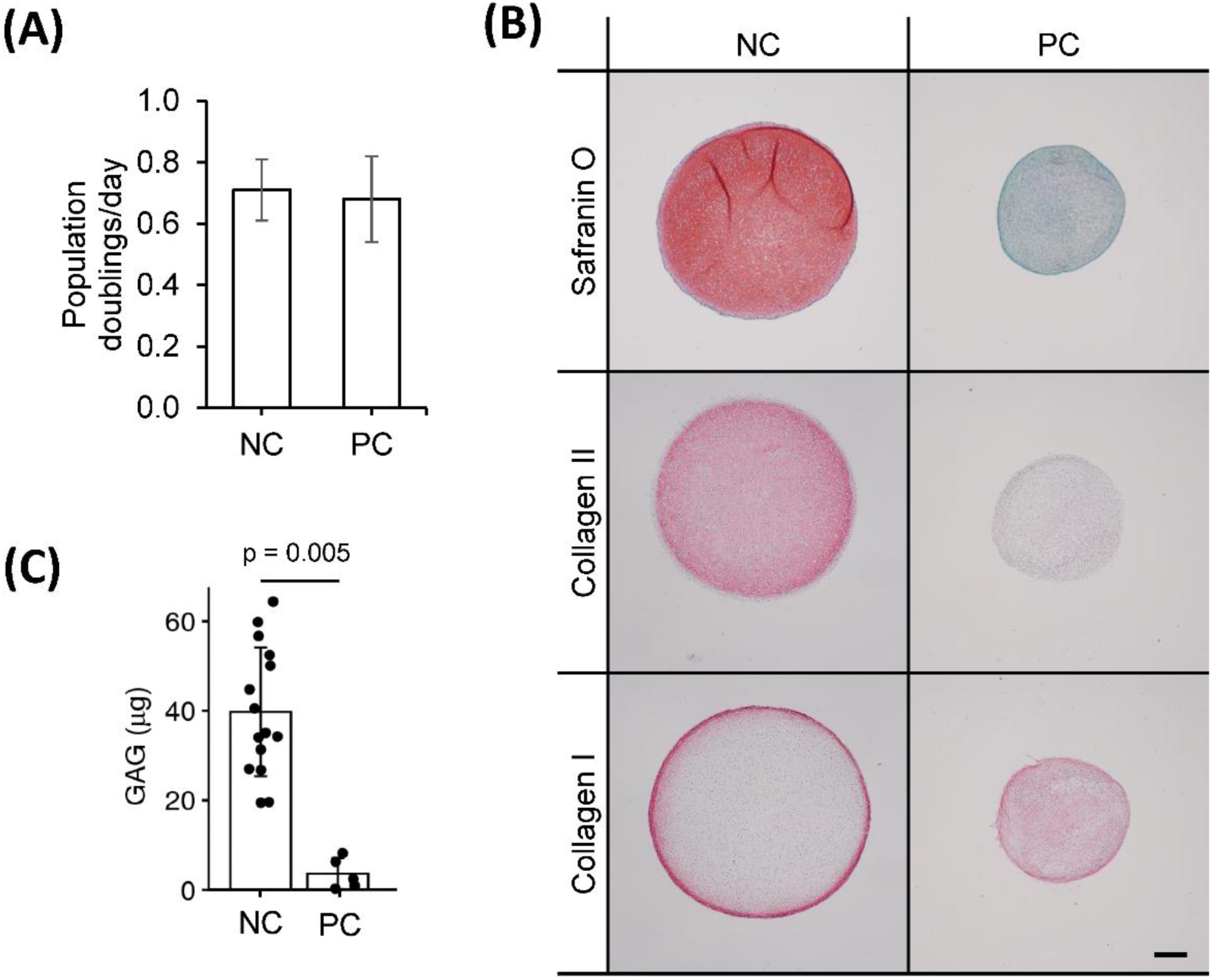
Chondrogenic capacity of perichondrial cells. **(A)** Proliferation rates of nasal chondrocyte (NC) and perichondrial cells (PC). **(B)** Biochemical quantification of NC and PC chondrogenic pellets. T-test p-value displayed. **(C)** Safranin O staining and immunohistochemical staining of pellets engineered from nasal chondrocytes (NC) and perichondrial cells (PC). Scale bar is 200 µm

### 3.3 Identity assay

We sought to distinguish the cells from these two tissues based on their gene expression profiles. NC cells expressed significantly higher levels of type II collagen and relative ratios of collagen II:I, aggrecan:versican and, at passage two, HAPLN1:MFAP5; whereas PC cells expressed significantly higher levels of versican, MFAP5, and nestin (Fig 3). Expanded cells were then cultured as 3D micromass pellets in chondrogenic conditions for two more weeks. NC cells from engineered pellets expressed significantly more collagen II and higher ratios of collagen II:I, aggrecan:versican, and HAPLN1:MFAP5, and PC cells expressed significantly higher levels of versican and MFAP5 (Fig S1). In summary, these results demonstrate that nasal chondrocytes and perichondrial cells have statistically significant differential expression of cartilage-related genes both during expansion and after pellet culture.

**Figure 3.**
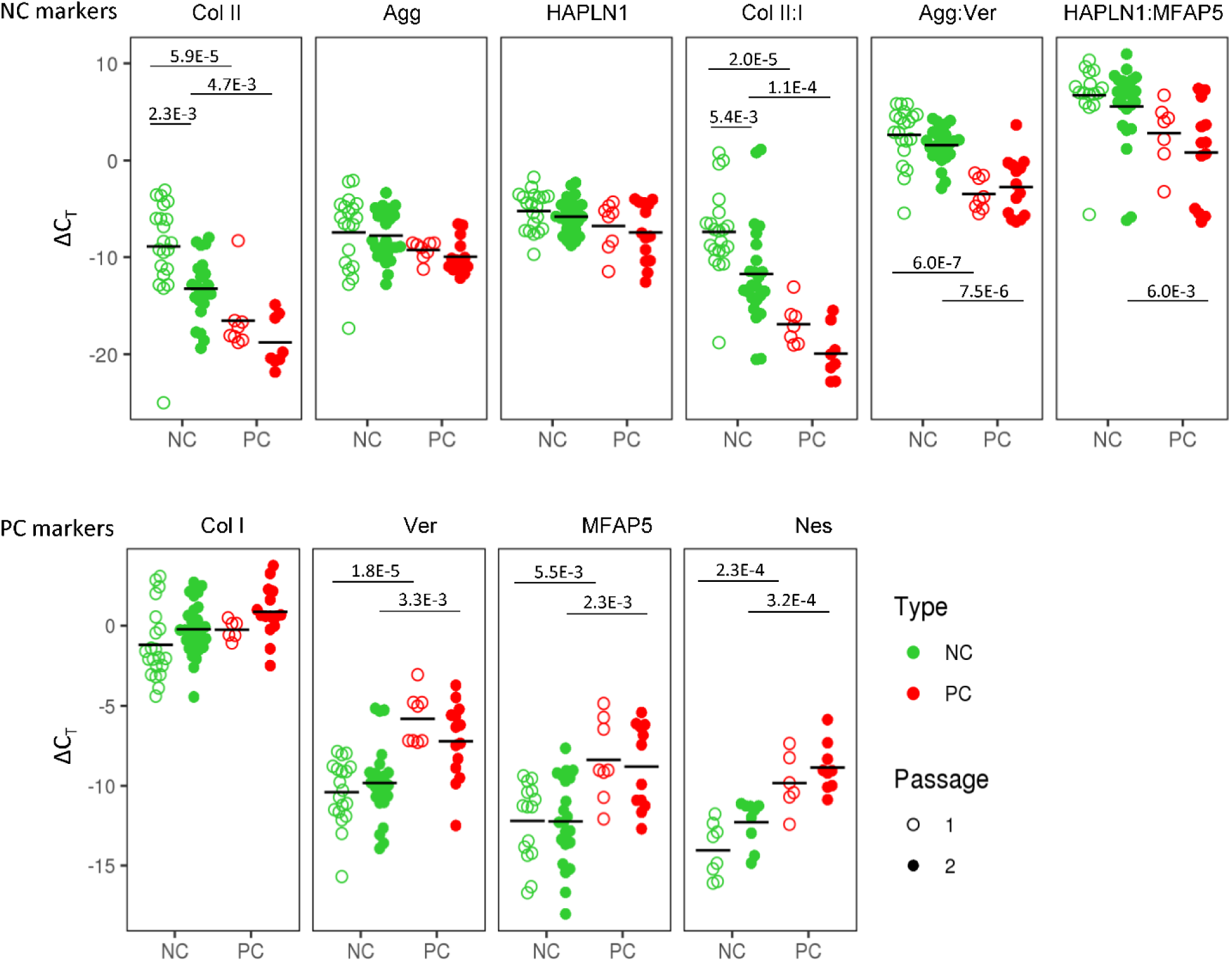
Nasal chondrocyte and perichondrial cell identity. Gene expression comparisons between pure nasal chondrocyte (NC) and pure perichondrial cell (PC) populations at passage one and two. The fold change expression relative to GAPDH is displayed. Bonferroni multiple comparison corrected p-values are displayed

### 3.4 Purity assay

The only method currently available to assess the purity of the starting native cartilage biopsy is by manually counting the number of cells in each type of tissue in a histological image (Fig S2). This method suffers from limitations due to histological artifacts, unclear distinction between tissue types, its semi-quantitative and destructive nature, and the fact that a histological section may not be representative of the whole tissue. Here we assessed if the purity of a *mixed* cell population could also be estimated based on gene expression analysis.

Spearman correlation coefficients (ρ) of the gene expression of cells at passage two that we combined at specific ratios of NC and PC cells revealed statistically significant trends across donors. Due to high donor-to-donor variability, the correlations between cell population purity and gene expression were higher per donor per gene than across donors. The highest correlation was found for the relative expression of aggrecan:versican (ρ = 0.69), where the ratio was higher in purer populations containing more NCs; per donor the correlations were even stronger (ρ = 0.61-0.98) (Fig S3A).

In general, more significant differences in gene expression in individual genes and cell purity were seen at passage two compared to the pelleted cells’ gene expression (Fig S3B). Therefore, we focused on passage two for the subsequent purity model.

We performed multiple logistic regression to compare gene expression of collagen type I and II, aggrecan, versican, MFAP5, HAPL1, and nestin to the cell population purity.To gain insight into which genes were most important, stepwise selection (Agostini et al., 2015) was implemented and the model with the lowest Akaike information criterion (AIC) was chosen. Versican and collagen type II were found to be the factors most predictive of purity and significantly contributed to the model (p-value = 2.7e-3, p = 2.8e-3, respectively, and overall, the model was significant (Hosmer-Lemeshow p = 0.95 and McFadden pseudo R^2^ = 0.53). The coefficient estimates from the model and the ΔCt values for versican and collagen II can be used to estimate the purity of a population of nasal cartilage-derived cells using Equation 1, where inverse logit is exp(x)/(1+exp(x)).

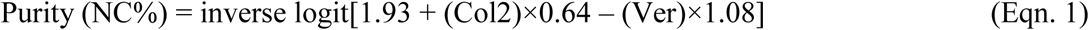

The purity predicted by the model was plotted against the known purity and the resulting R^2^ value of the observed and predicted values was 0.79 (Fig 4B).

**Figure 4.**
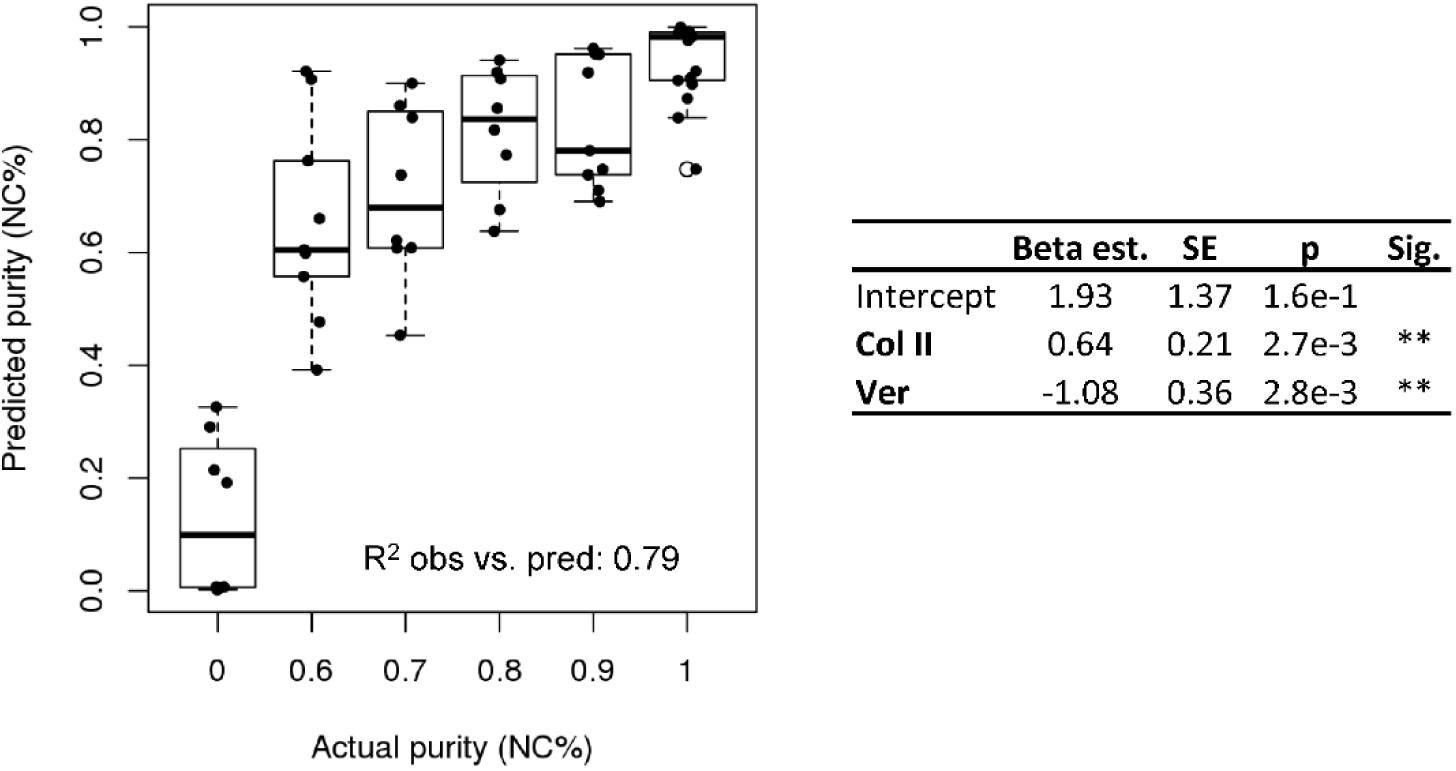
Purity assay. Purity assay results. Multiple logistic regression model based on the expression of collagen II and versican to estimate cell purity (NC%)

### 3.5 Potency assay

We investigated whether predictive gene expression markers can be used to estimate the capacity of the cells to form engineered cartilage. The final cartilage quality is currently assessed using the Modified Bern Score (MBS), a semi-quantitative score of safranin O-stained histological images (Lehoczky et al., 2019; Grogan et al., 2006), and via GAG quantification (Thej et al., 2018).

GAG content as well as the histological MBS score of the chondrogenic pellets were positively correlated to the cartilage identity gene expression markers and negatively correlated to the perichondrial identity markers (Fig S4). The interrelationship of potency and purity is visualized in the top left corner of the correlation plot, which shows that purity (NC%), GAG, GAG/DNA, and MBS are highly correlated (Fig 5).

**Figure 5.**
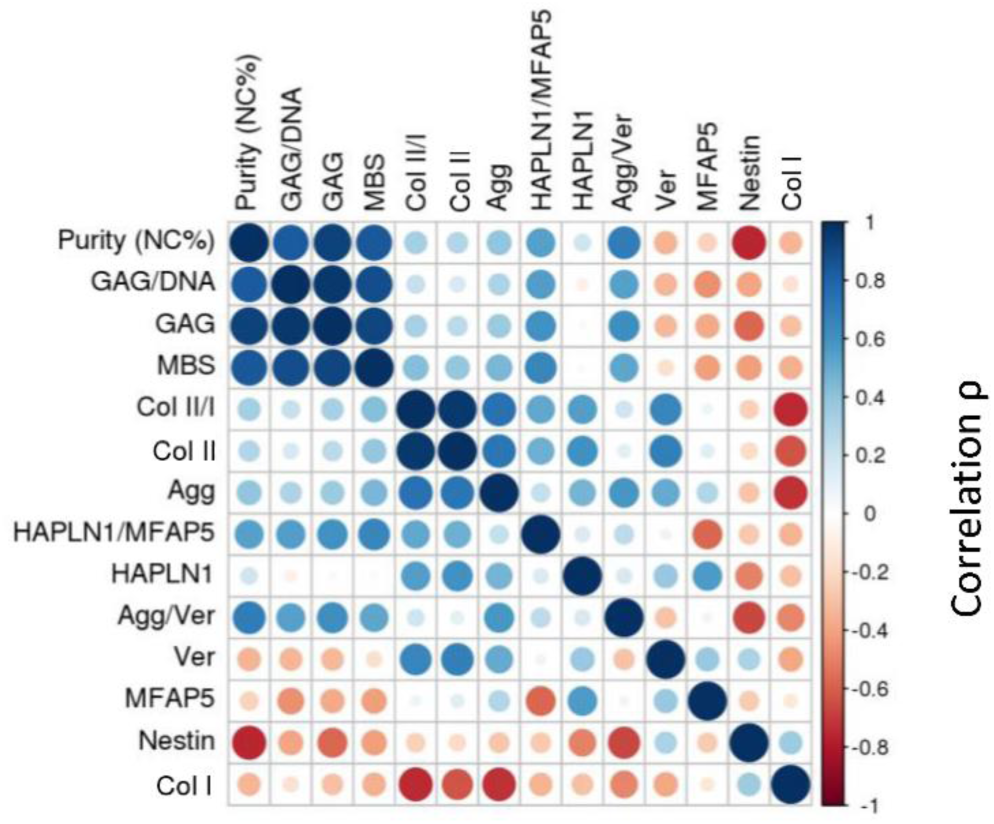
Correlation plot with passage two gene expression. Spearman correlations (ρ) depicted for passage two gene expression

More significant gene expression trends were seen when analyzing the cells at passage two compared to after engineered pellet culture, so we developed a potency assay for this time point.

In order to develop a potency assay that could predict the amount of GAG in the final engineered cartilage based on the gene expression of the starting cell population, we trained a generalized linear model with a log-link and gamma distribution. The gamma distribution was selected because it only predicts positive values and because its distribution is flexible enough to fit many response shapes (Hardin & Hilbe 2007). To select which gene expressions could best predict GAG produced by cells culture as pellets, stepwise selection was performed. Collagen II and MFAP5 were found to be the most significant and the model showed good results (training R^2^ = 0.34 and testing R^2^ = 0.78 of observed vs. predicted values; Figure 6A). The equation of the potency assay to predict the amount of GAG produced via the gene expression of passage two cells (Equation 2), where the ΔCt values of the genes should be entered, was generated using both the test and training data together, to report the most accurate coefficient estimates possible.

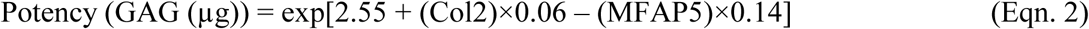

**Figure 6.**
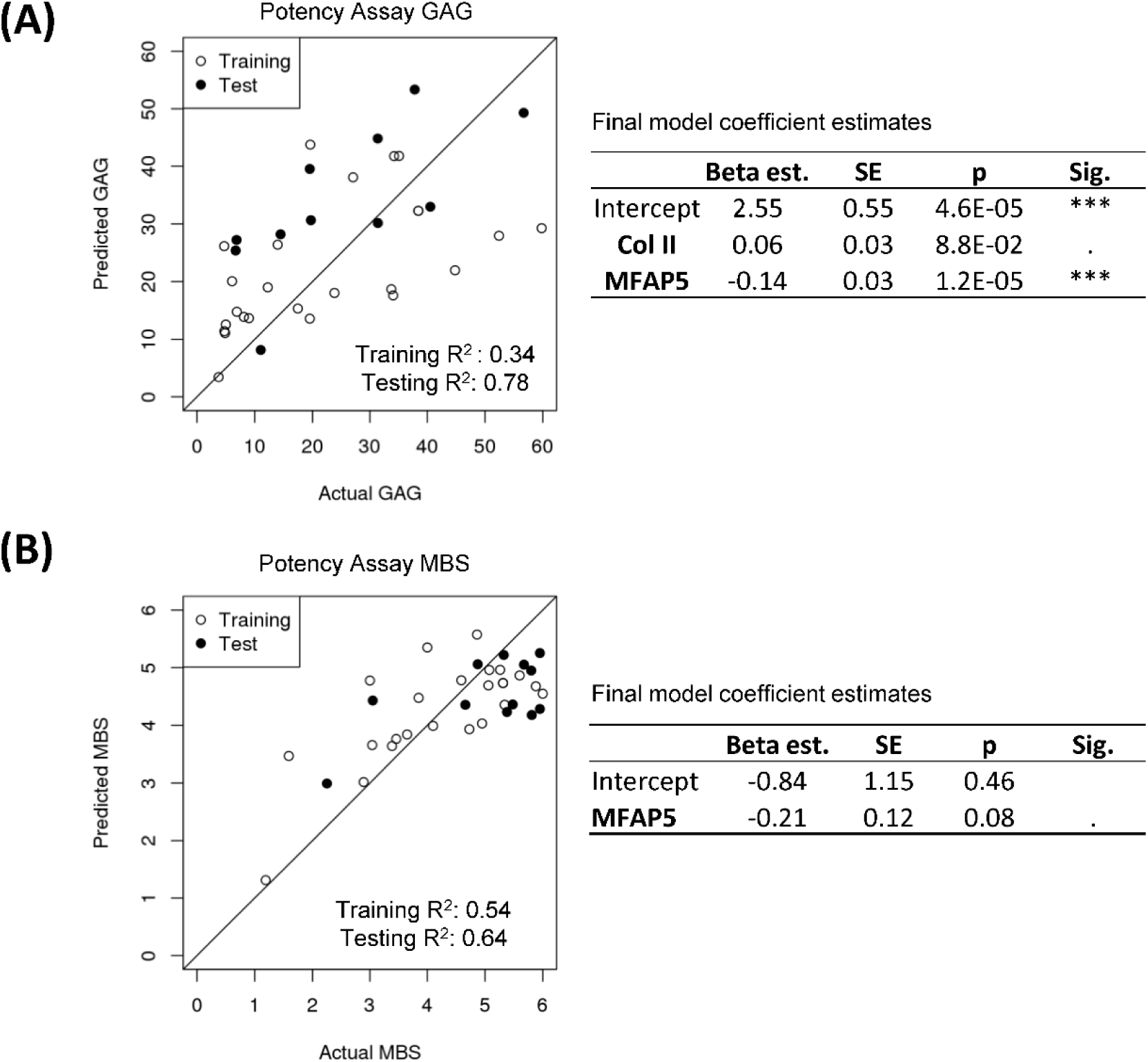
Potency assay. **(A)** Generalized linear model with a gamma distribution and log-link to predict GAG. The estimated model coefficients, standard errors (SE), and significances are calculated with the training and test data combined. **(B)** Multiple logistic regression model to predict MBS. The estimated model coefficients, standard errors (SE), and significances are calculated with the training and test data combined

For a potency assay that can predict the histological score of the final engineered cartilage from passage two gene expression, a logistic regression model was trained. Stepwise selection found that the best model included only the MFAP5 gene, and showed good predictive ability in this dataset (training R^2^ = 0.54 and testing R^2^ = 0.64 of observed vs. predicted values; Figure 6B). The equation of the potency assay to predict the histological MBS score (Equation 3), where the ΔCt value of the gene should be used, was generated with both the training and test data together. Again, the inverse logit is exp(x)/(1+exp(x)), imposing upper and lower bounds on the model.

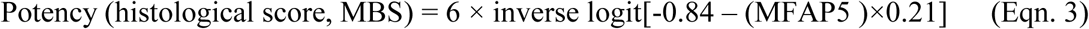

### 3.6 Implementation of in-process controls

Since isolated nasal septum-derived cell populations may include some perichondrial cells, we tested the impact of various amounts of contaminating cells on final engineered cartilage quality. The chondrogenic capacity of contaminated cell populations was consistently lower than of pure cell populations, as observed across 15 donors, demonstrated by safranin O staining, GAG quantification, and immunohistochemical analysis of collagen types II and I (S5). We confirmed the negative effect of perichondrial cells on the engineered cartilage not only in pellet culture, but also when produced according to the clinical trial protocol where cells are seeded onto a collagen I/III scaffold (Fig S6).

A threshold of acceptable purity needs to be set to guarantee the quality of the final product. Known quantities of NC and PC cells were mixed together and chondrogenic pellets were produced. Histological scoring was then used to set acceptable limits of PC cell contamination so that the quality of the final product would still meet the clinical trial release criteria (MBS ≥ 3). Due to donor-to-donor variability, potential cross-contamination from the mechanical tissue separation method, and considering the limitations of histological analysis, we show that some donors could still produce cartilage matrix of sufficient quality with up to 40% PC contamination, while the less potent donors could produce cartilage matrix with a PC contamination of up to 30% PC cells (Fig 7A).

**Figure 7.**
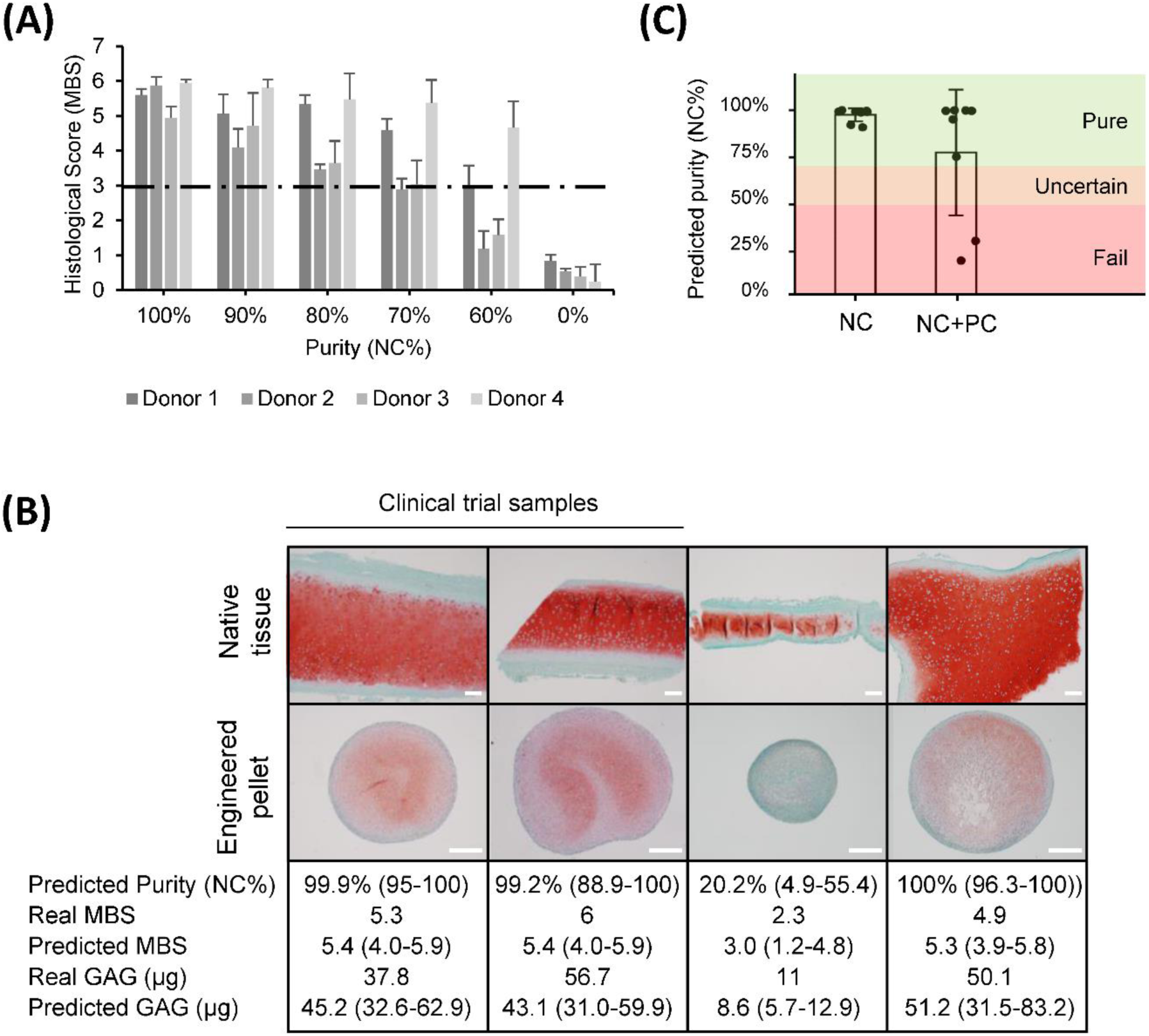
Quality estimation and in-process control implementation. **(A)** Histological scores (Modified Bern Score) of engineered pellets derived from specific starting population purities. **(B)** Safranin O stained native and engineered cartilage harvested for a clinical trial study or for other purposes. The predicted purity and 95% CI, real MBS, predicted MBS and 95% CI, real amount of GAG, and predicted GAG with 95% CI. Two of the samples were produced from samples deriving from clinical trials. Scale bars are 200 µm. **(C)** Predicted purities of pure (NC) and mixed (NC+PC) biopsies

Using the purity and potency assays we developed, the quality was estimated based on the passage two gene expression of cell populations for clinical trial samples and for heterogeneous biopsies collected from patients that underwent plastic surgeries with variable amounts of overlaying perichondrium. The predicted histological score results closely matched the actual values, and the quantified amounts of GAG could be estimated well, predicting if cells would produce high or low amounts of GAG (Fig 7B). The clinical trial starting materials were assessed to be pure and the potency assays predicted good chondrogenic capacity which was confirmed by the high quality of the engineered cartilage. The more heterogeneous cartilage samples from plastic surgery procedures had more variable results. The purity assay predicted the worst sample to have a purity of 20%, many samples to be 99% pure, and the mean purity of mixed samples to be 75% (Fig 7C). Consistent with the established purity threshold, cells that were predicted to be >70% pure were all able to produce cartilaginous tissues that passed the histological score release criteria. The sample with a predicted purity of 20%, on the other hand, produced a pellet that failed the release criteria (histological score = 2.3).

## 4 Discussion

In this study, we established novel, in-process controls to ensure the quality and standardization of nasal chondrocyte-based engineered cartilage grafts. Histological analysis revealed that nasal septal cartilage may be harvested with some adjacent tissue, and that there may still be fragments of perichondrial tissue overlaying the cartilage even after a trained operator further separates the tissues. Although some researchers claim that perichondrial cells from other cartilage sources have chondrogenic potential (Hellingman et al., 2011), we discovered that unlike chondrocytes, nasal septal perichondrial cells do not have the capacity to form GAG- and collagen type II-rich engineered tissues. We found that increasing amounts of perichondrium in the starting material profoundly decreases the quality of engineered cartilage, as seen by less GAG and collagen II production during chondrogenic culture. Therefore, minimal contamination of perichondrial cells must be ensured. The NC identity marker we found is collagen II, and the PC identity markers were, versican, MFAP5, and nestin. To quantitatively determine the percentage of contaminating cells in a population, we developed a model that correlates the expression of multiple gene expression markers to the purity of a cell population. Similarly, to predict the chondrogenic capacity of a cell population, we built models to estimate GAG production and the final histological MBS score in engineered cartilage. Finally, we discuss how such quality controls could be implemented during the production of cell or tissue therapies.

In practice, polymerase chain reaction (PCR) instrumentation is ubiquitous, so a gene expression-based quality control could be easily implemented. The cost of the quality control assay could be reduced by selecting a handful of genes for a standard qPCR analysis compared to transcriptomic analysis or single-cell RNA sequencing, and for a routine test could be enough information to confirm cell identity (Maertzdorf et al., 2016).

To implement such gene expression-based quality controls, a suitable time point during the manufacturing process must be chosen. Biomarkers vary not only spatially within the tissue, but also temporally during monolayer expansion and after tissues are engineered (Detela et al., 2019; Späth et al., 2018; Tay et al., 2004), and it also may be that the cells have more distinct gene expression profiles at certain time points than others (Tekari et al., 2014). From a practical perspective, an earlier quality control would save costs, because the quality of the cells could be established before an expensive production is undertaken. However, after one or two weeks of cell expansion, there are many more cells and an aliquot can be taken without depleting the whole cell population and the gene expression analysis of an aliquot of a cell suspension provides a broad readout of the total cellular material. Obtaining cells before they are embedded in the scaffold would allow to perform the analysis nondestructively. Importantly, the biomarkers we investigated had the most distinct expression levels after the expansion phase. Consequently, we propose that our gene expression-based assays should be implemented on expanded passage two cells.

Generalized linear models for the development of gene expression-based quality controls for regenerative medicine is a natural extension of their use in biomarker-based disease diagnosis (Hosmer et al., 2013; Faraway et al., 2006). Here we show how multiple logistic regression can be used to model purity percentages with the advantage of being able to provide biologically relevant estimates, i.e., between 0 and 100% (Zhao et al., 2001). When further screening the most significant genes that contributed to the purity model with stepwise selection (Liu et al., 2019; Ying et al., 2018), we found the combination of collagen II and versican expression to be predictive, a relatively uncommon gene pair compared to the often studied gene expression ratios of collagen II:I and aggrecan:versican. The selection appears reasonable, with one chondrocyte marker, collagen II, and one perichondrium marker, versican, used in the model and being inversely related to each other.

We showed how logistic regression can be used to estimate the histological score, a value bounded between 0 (worst) and 6 (best). Estimating GAG required modeling positive values only, therefore, we demonstrated how a generalized linear model with a gamma distribution and log-link could be implemented, similarly to other biomarker applications (Schufreider et al., 2015; García-Broncano et al., 2014). Stepwise selection was used again for the potency models, returning the combination of collagen II and MFAP5 for prediction of GAG, and MFAP5 alone for modeling histological MBS score. Increased MFAP5 expression has been correlated with decreased chondrogenic potential in mesenchymal stem cells (Solchaga et al., 2010). MFAP-5 protein binds active TGFβ1, TGFβ2, and BMP2, sequestering these pro-chondrogenic factors in the matrix (Combs et al., 2013). Intracellular MFAP5 has been shown to bind and activate notch signaling (Miyamoto et al., 2006), which inhibits the regulator of cartilage formation, Sox9 (Hardingham et al., 2006). Notch has been found primarily in the perichondrium rather than the cartilage layer in mandibular condylar cartilage (Serrano et al., 2014), which, like nasal cartilage, is derived from cranial neural-crest cells (Chai et al., 2000). The predictive ability of these models are significant especially when considering that only ∼30-40% of the variance in protein abundance is explained by mRNA levels (Vogel et al., 2012). The selection of different genes for each potency assay may be due to the fact that they assess quality in slightly different ways; the histological score includes information not only of the GAG content, but also about the morphology of the cells.

We observed that all pellets that contained at least 70% NC cells pass the clinical trial release criteria, i.e., histological score ≥ 3, but more contamination could also lead to good results in some cases. To implement the purity assay, we propose a conservative three-category rating scale for the predicted purity (NC%), i.e., if cells are estimated to be more than 70% pure, they are labeled as *pure*, less than 50% pure, they are labeled as *fail*, otherwise the estimation is labeled as *uncertain*. We propose to introduce this uncertain region for the time being until further data can be collected and the estimates can be made more precise. In practice, we would recommend that starting cell populations labeled as pure or uncertain should continue in the production process, however, if the cells fail the purity test, the costly production should be halted. Cartilage engineered from cells of uncertain purity would nevertheless need to pass the release criteria (such as the histological score-based release criterial), ensuring the quality of the product.

The selected genes and coefficient estimates for the models will have to be updated as more data are obtained, and the in-process controls will have to be validated to meet GMP standards. Moreover, the definition of a high quality graft may need to be revised as more long-term clinical outcome data are collected.

In conclusion, we have put forward gene expression-based assays for identity, purity, and potency to help ensure the safe and effective clinical use of nasal chondrocyte-derived engineered cartilage. More generally, we provide an example of the development and implementation of purity and potency assays based on relatively simple qPCR assays, stepwise selection of the most significant genes, and predictive in silico models. This approach could be relevant for the development of quality controls for other products in the emerging field of regenerative medicine, one of the biggest challenges for advanced therapy medicinal products to overcome for clinical translation.

## 6 Author Contributions

AA conceived and designed the study. AA and LP performed the experiments, analyzed the data and wrote the manuscript. LP created the models. AB contributed to the study design and revised the manuscript. MH and RK contributed to the sample preparation. IM contributed to compiling the data and critically revised the manuscript.

## 8 Acknowledgments

We would like to thank Sandra Feliciano for her help with sample analysis. Thank you to Dr. Florian Geier and Dr. Julien Roux for the statistical analysis discussions. Thank you Dr. Sylvie Miot and Anke Wixmerten for critically reading the manuscript.

