## Supplementary Information for "Biomarker signatures of quality for nasal chondrocyte-derived engineered cartilage"

### *Supplementary Material*

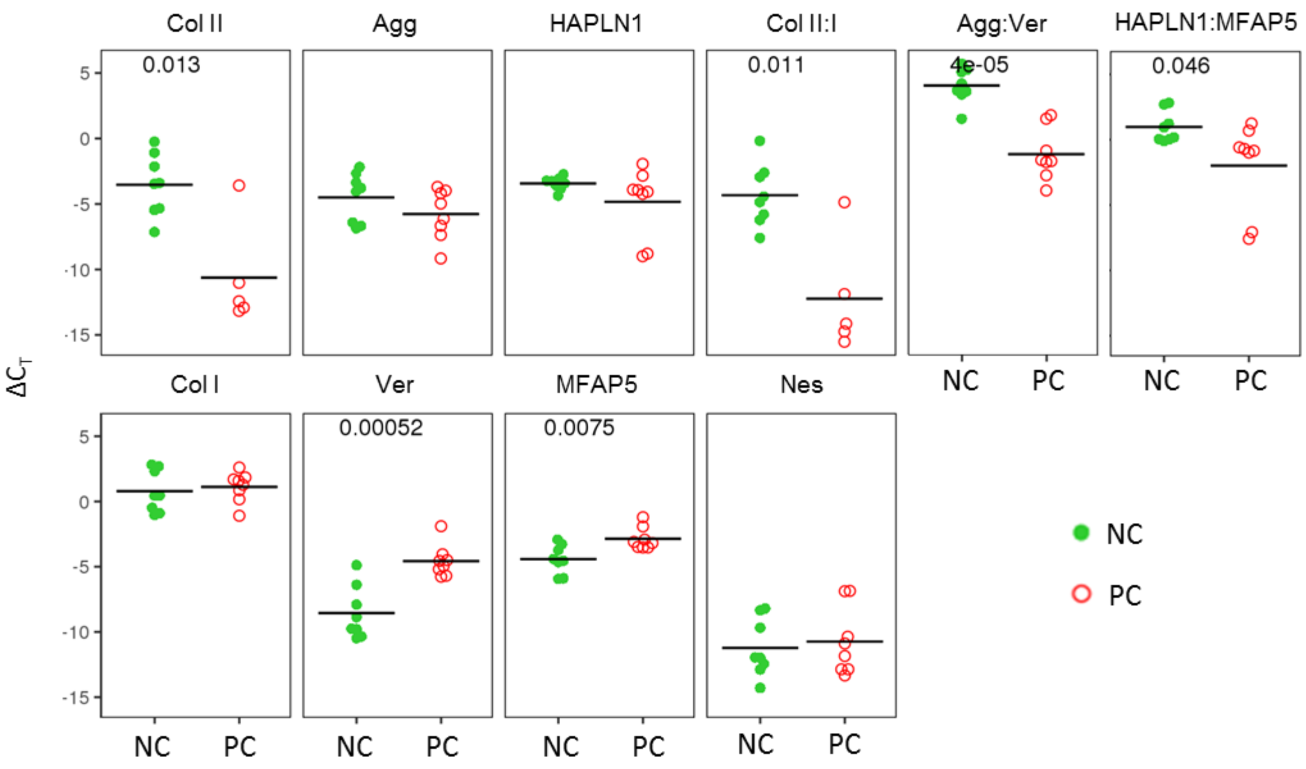


**Supplementary Figure 1.** Gene expression of cells after chondrogenic pellet culture. Gene expression of nasal chondrocytes (NC) and perichondrial cells (PC) after chondrogenic pellet culture. T-test p-values are displayed. Black lines indicate the mean values

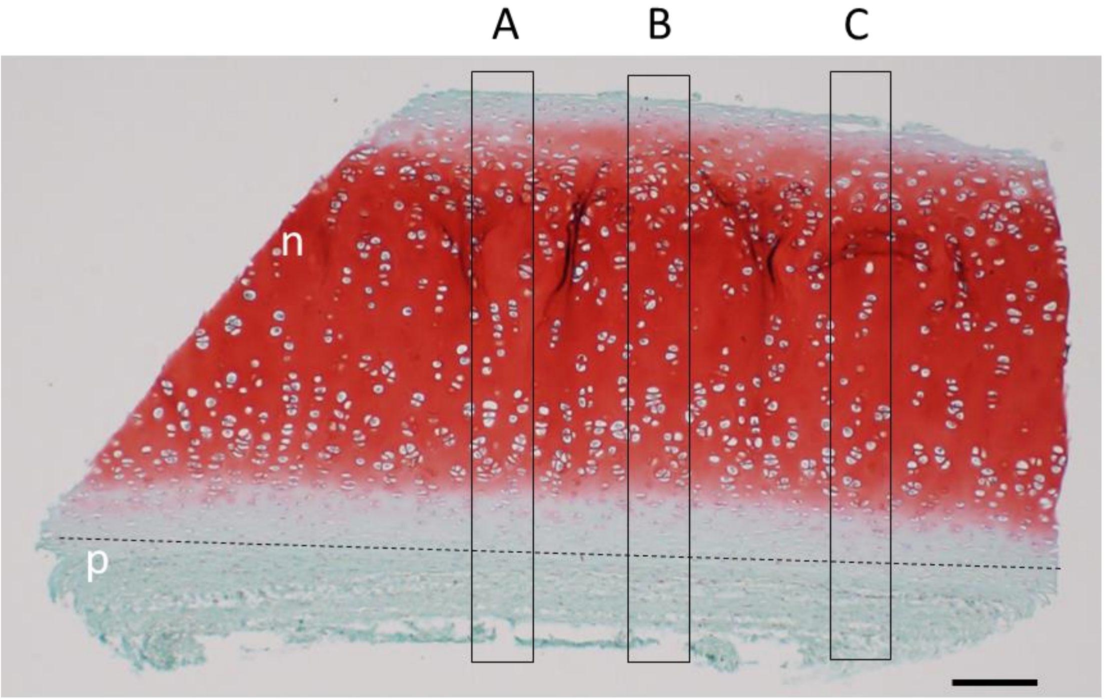


| | A | B | C | Average |
|---|---|---|---|---|
| **Num. NC** | 113 | 115 | 91 | |
| **Num. PC** | 11 | 15 | 17 | |
| **%PC** | 9.8 % | 13.0 % | 18 % | **13.6 %** |

**Supplementary Figure 2.** Histological assessment of native biopsy purity. Histological assessment of native cartilage biopsy purity is done by taking three complete cross sections (A-C), labeling the nasal cartilage and perichondrium based on the structure of the tissues, with cartilage having lacuna in which cell reside and perichondrium having a more fibrotic texture, and manually counting the number of nuclei in each part. Scale bar is 200 µm

**(A)** P2

| Replicates | Col II | Agg | HAPLN1 | Col II:I | Agg:Ver | HAPLN1: MFAP5 | Col I | Ver | MFAP5 | Nes |
|---|---|---|---|---|---|---|---|---|---|---|
| All, n = 48 | 0.35 | 0.38 | 0.32 | 0.38 | 0.69 | 0.50 | -0.23 | -0.26 | -0.28 | -0.51 |
| p | 7.1E-03 | 2.0E-03 | 1.1E-02 | 3.0E-03 | 7.9E-10 | 3.8E-05 | 7.2E-02 | 3.8E-02 | 3.0E-02 | 2.4E-05 |
| Donor 1 | 0.37 | 0.21 | 0.54 | -0.05 | 0.61 | 0.57 | 0.13 | -0.32 | 0.00 | -0.43 |
| Donor 2 | 0.40 | 0.52 | 0.42 | 0.90 | 0.98 | 0.96 | -0.32 | -0.68 | -0.91 | -0.59 |
| Donor 3 | 0.33 | 0.47 | -0.58 | 0.58 | 0.93 | 0.31 | -0.89 | -0.95 | -0.92 | -0.85 |
| Donor 4 | 0.88 | 0.81 | 0.61 | 0.94 | 0.84 | -0.10 | -0.3 | -0.11 | 0.57 | -0.83 |

ρ 1 0 -1

**(B)** Pellets

| Replicates | HAPLN1:MFAP5 | Agg:Ver | Ver | MFAP5 |
|---|---|---|---|---|
| All, n = 48 | 0.49 | 0.67 | -0.68 | -0.43 |
| p | 1.5E-02 | 3.0E-04 | 3.0E-04 | 3.8E-02 |
| Donor 1 | 0.94 | 0.83 | -0.83 | -0.77 |
| Donor 2 | 0.60 | 0.77 | -1.00 | 0.09 |
| Donor 3 | 0.94 | 0.94 | -0.77 | 0.14 |
| Donor 4 | 0.43 | 1.00 | -0.94 | -0.94 |

**Supplementary Figure 3.** Correlations between purity and gene expression levels. Spearman correlation coefficients (ρ) of purity (NC%) vs. gene expression for cells **(A)** expanded to passage two and **(B)** after chondrogenic pellet culture from four donors and 48 experimental replicates. First the correlation coefficient across all independent experiments is displayed with the p-value, then per individual donor correlations (ρ)

**(A)**

P2

GAG

| Replicates | Col II | Col II:I | Agg:Ver | HAPLN1: MFAP5 | MFAP5 | Ver |
|---|---|---|---|---|---|---|
| All, n = 46 | 0.35 | 0.47 | 0.46 | 0.59 | -0.50 | -0.49 |
| p | 2.3E-02 | 1.0E-03 | 2.0E-03 | 1.9E-05 | 4.4E-04 | 1.0E-03 |
| Donor 1 | 0.70 | 0.31 | 0.94 | 0.83 | -0.37 | -0.77 |
| Donor 2 | 0.49 | 0.94 | 1.00 | 1.00 | -0.94 | -1.00 |
| Donor 3 | 0.49 | 0.60 | 0.83 | 0.20 | -0.94 | -0.83 |
| Donor 4 | 0.94 | 1.00 | 0.83 | 0.09 | 0.60 | -0.26 |

MBS

| Replicates | Col II | Col II:I | Agg:Ver | HAPLN1: MFAP5 | MFAP5 | Ver |
|---|---|---|---|---|---|---|
| All, n = 40 | 0.44 | 0.48 | 0.45 | 0.65 | -0.49 | -0.30 |
| p | 5.0E-03 | 2.0E-03 | 4.0E-03 | 6.0E-06 | 1.0E-03 | 5.9E-01 |
| Donor 1 | 0.55 | 0.14 | 0.89 | 0.77 | -0.26 | -0.89 |
| Donor 2 | 0.49 | 0.94 | 1.00 | 1.00 | -0.94 | -1.00 |
| Donor 3 | 0.77 | 0.83 | 1.00 | 0.49 | -0.94 | -1.00 |
| Donor 4 | 0.94 | 1.00 | 0.83 | 0.09 | 0.60 | -0.26 |

ρ: 1, 0, -1

**(B)**

Pellets

GAG

| Replicates | Agg:Ver | Ver |
|---|---|---|
| All, n = 46 | 0.54 | -0.54 |
| p | 3.6E-04 | 4.6E-04 |
| Donor 1 | 0.83 | -0.83 |
| Donor 2 | 0.77 | -1.00 |
| Donor 3 | 0.77 | -0.49 |
| Donor 4 | 1.00 | -0.94 |

MBS

| Replicates | HAPLN1 | Agg:Ver | HAPLN1: MFAP5 | Ver | MFAP5 |
|---|---|---|---|---|---|
| All, n = 40 | 0.35 | 0.43 | 0.49 | -0.49 | -0.42 |
| p | 0.048 | 0.013 | 0.004 | 0.004 | 0.016 |
| Donor 1 | 0.09 | 0.71 | 0.89 | -0.94 | -0.83 |
| Donor 2 | 0.83 | 0.77 | 0.6 | -1 | 0.09 |
| Donor 3 | 0.6 | 0.94 | 0.94 | -0.77 | 0.14 |
| Donor 4 | -0.6 | 1 | 0.43 | -0.94 | -0.94 |

**Supplementary Figure 4.** Correlations between quality and gene expression levels. Spearman correlation coefficients (ρ) of GAG (from 15 donors and 46 experimental replicates) and the histological score (Modified Bern Score; MBS; from 15 donors and 40 experimental replicates) vs. gene expression for cells **(A)** expanded to passage 2 and **(B)** after chondrogenic pellet culture. First the correlation coefficient across all independent experiments with the p-value are displayed, then per individual donor correlations (ρ)

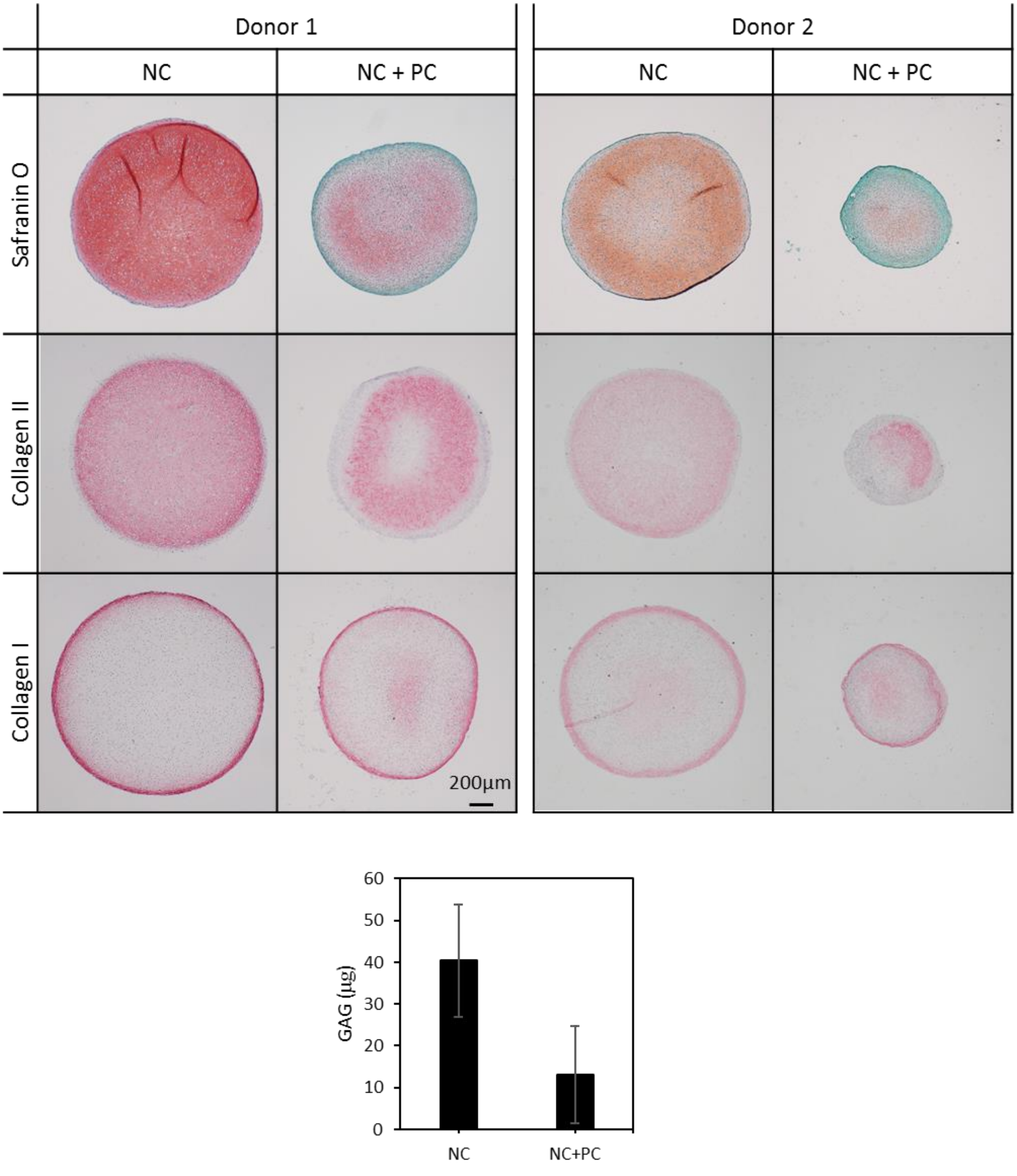


**Supplementary Figure 5.** Engineered cartilage pellets from pure (NC) and contaminated (NC+PC) cell populations. Safranin O staining, immunohistochemical staining for collagen II and I, and biochemically quantified GAG (µg) of pellets engineered from cells that were isolated from pure or contaminated nasal septal biopsies from 15 donors

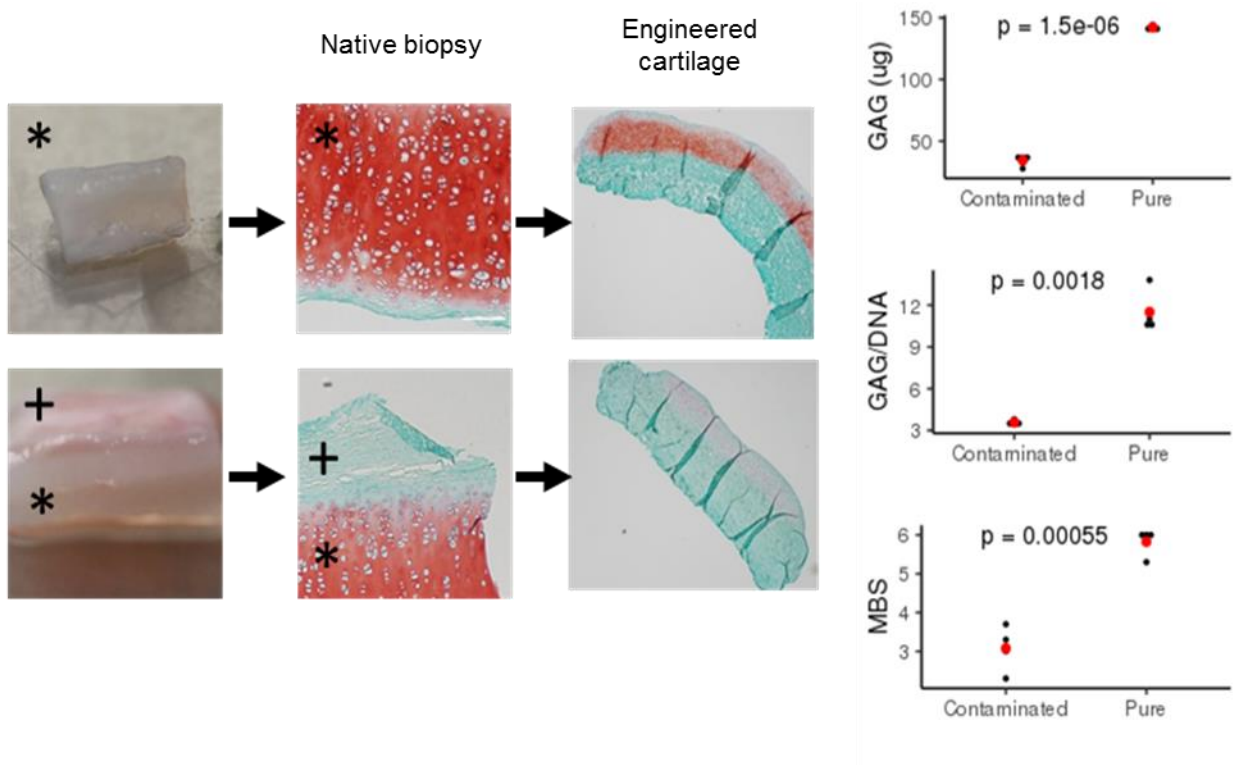


**Supplementary Figure 6.** Cartilage engineered on scaffolds from pure and contaminated biopsies. Safranin O staining and biochemical quantification of tissues engineered on collagen I/III scaffolds with cells from a pure cartilage biopsy or a cartilage biopsy contaminated by perichondrium. The mean is displayed with a red dot. Symbols are * nasal cartilage and + perichondrium
